# Not all roads lead to selfing: mating-system stability in invasive ectomycorrhizal fungi

**DOI:** 10.64898/2026.09.19.752884

**Authors:** Yi-Hong Ke, Yen-Wen Wang, Anna L. Bazzicalupo, Lotus Lofgren, Sara Branco, Rytas Vilgalys

## Abstract

Baker’s rule predicts that selfing is a beneficial trait for colonizers, whose prediction in the kingdom Fungi is reinforced by recent work showing the invasive ectomycorrhizal basidiomycete *Amanita phalloides* with self-incompatibility (SI) system has evolved unisexual reproduction in its introduced range. Unlike in plants, the multiallelic SI system of basidiomycetes does not fully prevent selfing among the meiotic offspring of a single dikaryotic (diploid) parent. This distinction allows observation of an intermediate state of selection where selfing is preferred in a species with SI system during colonization before a change in self-incompatibility architecture. Here, we analyzed the genomes of 189 isolates of the ectomycorrhizal basidiomycete *Suillus luteus* collected from both its native European range and five independently introduced regions to test whether selfing rates increased and whether the SI system had degraded after introduction. Unlike the *A. phalloides* case, we found no evidence of unisexual reproduction or loss of the SI system in the introduced ranges of *S. luteus*. Rather, all introduced populations maintained outcrossing rates higher than expected under random mating as in the native population. However, selfing events were detected in three individuals from a single introduced population, representing the first documented case of selfed sporocarps of Boletales fungi. Our work in *S. luteus* demonstrates that although selfing is possible in this species, it is not favored during colonization, potentially due to inbreeding depression and higher population density, highlighting the complex interaction among invasion biology, genetics, and the colonizing context.

## Introduction

Baker’s rule predicts that self-compatible species have an advantage during colonization of new territory because they can reproduce when mates are scarce^1^. This prediction has received mixed support in plants and has rarely been tested in fungi^2–4^. The most systematic study of breeding system change in fungi was conducted in yeasts, and suggests that “all roads lead to selfing”^5,6^, highlighting the potential selection for selfing in the broader fungal evolution context. It was recently discovered that the invasive ectomycorrhizal basidiomycete *Amanita phalloides* can produce unisexual sporocarps in its introduced range^7,8^, which represents an extreme form of selfing, echoed the same principle and provided the first evidence in support of Baker’s rule in a filamentous fungus. However, whether this represents a general response or an exception remains unknown. Unlike the plant self-incompatibility (SI) system, which fully prevents self-fertilization, the multiallelic mating-type system of basidiomycetes only reduces the probability of mating between monokaryotic (haploid) siblings carrying the same mating type^9,10^, while selfing among other siblings from a single dikaryotic (diploid) parent remains possible. Investigating a fungus that recently expanded to new territories offers a unique opportunity to test selection on selfing in a species with an SI system during colonization, independent of SI system breakdown. Here, we used the whole-genome sequences of 189 isolates of the basidiomycete *S. luteus*, spanning the native Central European range of this species, along with five regions where it has been independently introduced (South America, North America, New Zealand, Australia, and Africa) to test whether selfing rates increased and whether the SI system had degraded after introduction.

## Results and Discussion

### Self-incompatibility architecture remains intact in introduced populations

We found that all sampled individuals from both native and introduced populations were dikaryotic and contained two distinct alleles of *HD MAT*, the SI locus of basidiomycete. By assembling the highly divergent alleles of the mating-type-determining *HD MAT* locus, which contains the *HD1* and *HD2* genes^11^, we recovered two heterogeneous alleles from each genome, whose pairwise nucleotide identity was always less than 83.7%. Pairwise identities of the protein sequences of the N-terminal regions of the two *HD* proteins, the region directly involved in mating specificity, formed a disjunct distribution without intermediate values, where alleles either had > 98% or < 85% identity (Figure S1; Tables S1-S2). We therefore assigned alleles with > 98% identity as the same mating type and the rest as distinct mating types (Table S3). The same *HD1* mating type was always associated with the same *HD2* mating type without evidence of intralocus recombination, resulting in 66 distinct mating types among 378 alleles. No fruiting body containing a single mating type was detected in either native or introduced populations, indicating that the unisexual reproduction reported in invasive *A. phalloides*^7^, where sporocarps bypass *HD MAT* control by developing from a single haploid mycelium, is absent in *S. luteus*.

Despite their functional integrity, mating-type allele diversity declined in association with documented founder bottlenecks. At the genotype level, the 53 individuals from native Central Europe carried 53 unique genotypes without duplication, whereas all introduced populations contained duplicated genotypes despite smaller sample sizes (Table 1). Duplication was most extensive in Australia and South America, where the same genotypes occurred three or four times and multiple genotypes were sampled twice, matching the populations with the most severe documented genetic bottlenecks^12^. A similar pattern held at the allele level, where the native Central European population had the highest *HD MAT* allele diversity, reflected in the highest estimated total number of mating types based on the frequency of recurring alleles (Figure 1B; Table S4). Compared to the native population, all introduced populations contained a lower diversity of mating types and higher numbers of recurring mating types. The lowest allele diversity was again observed in the Australian and South American populations (Figure 1A). No mating types in one introduced population formed a subset of another (Figure S2), supporting independent introduction events from the diverse European source rather than sequential transfers between continents.

**Table 1.** *HD MAT* genotype duplication frequencies per population. Number of genotypes sampled in one, two, three, or four individuals.

| Population | Sample size | 1 time | 2 times | 3 times | 4 times |
| --- | --- | --- | --- | --- | --- |
| Central Europe | 53 | 53 | 0 | 0 | 0 |
| Africa | 1 | 1 | 0 | 0 | 0 |
| Australia | 32 | 16 | 3 | 2 | 1 |
| North America | 35 | 29 | 3 | 0 | 0 |
| New Zealand | 28 | 25 | 0 | 1 | 0 |
| South America | 40 | 25 | 6 | 1 | 0 |

**Figure 1.**
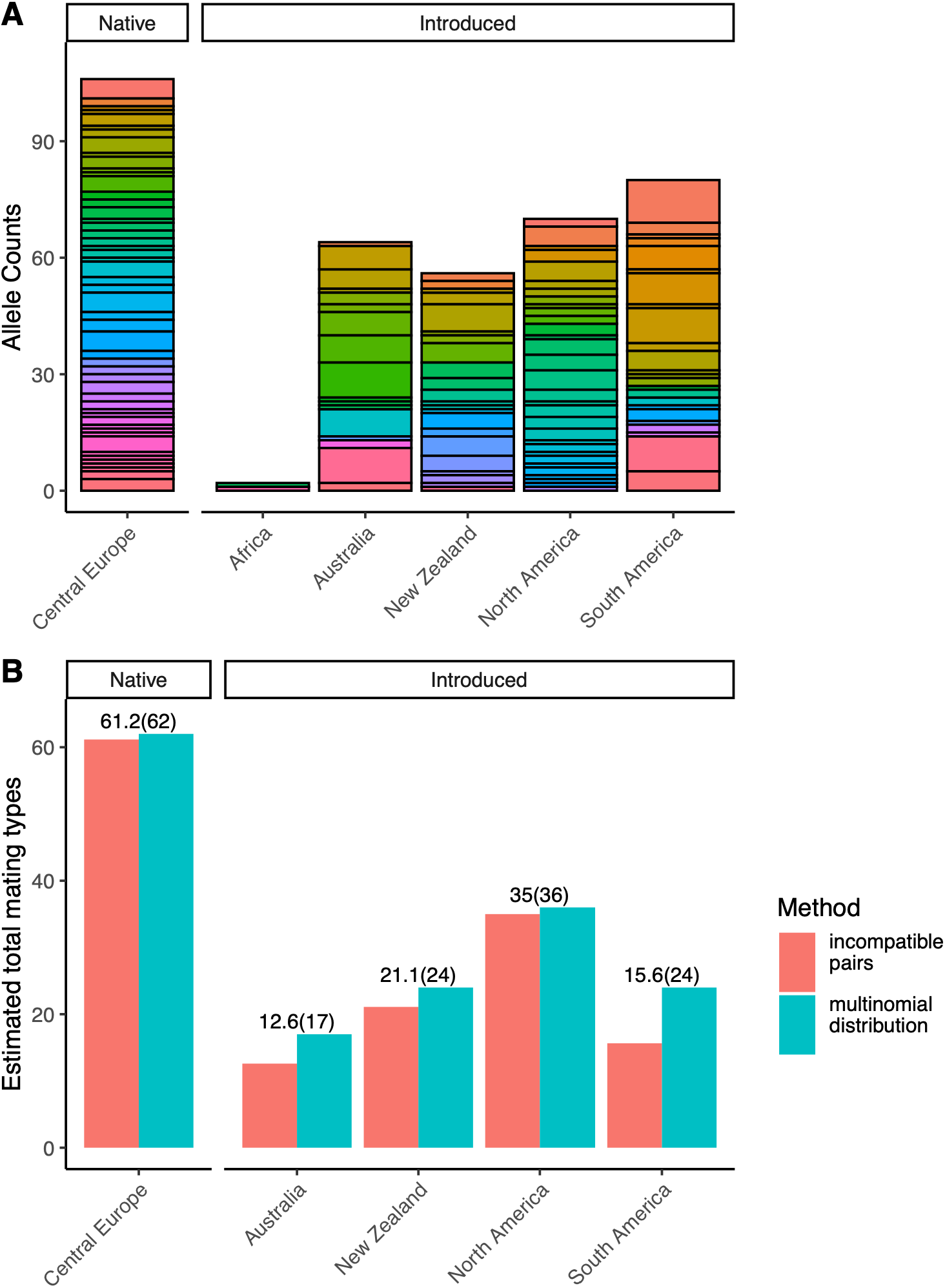
*HD MAT* allele diversity is reduced in introduced populations. (A) Counts of *HD MAT* alleles in each population of *S. luteus*. Each unique color within a stacked bar represents a distinct mating-type allele. Tall single-color segments in introduced populations indicate alleles sampled from multiple individuals. (B) Estimated total number of *HD MAT* alleles per population from incompatible-pair counts and maximum-likelihood estimation under the multinomial distribution.

### Outcrossing predominates at the population level

Despite the lower genetic diversity and a reduced number of mating types in the introduced *S. luteus* populations, the inbreeding coefficient (*F*_IS_) was negative across all populations (Table 2), indicating excess heterozygosity relative to the expectation under random mating (Hardy-Weinberg equilibrium). Notably, this excess heterozygosity is not unique to introduced populations of *S. luteus*, as it has also been documented in both native *S. luteus*, and other *Suillus* species^13–15^. The persistence of this signature in introduced populations indicates that the fundamental mating system has not shifted after introductions. To comply with the assumption of a panmictic deme, we limited the calculation to individuals collected in a single region or in nearby regions while retaining adequate sample size. We note that unaccounted population structure would only inflate the inbreeding coefficient via the Wahlund effect^16^, but not bias it toward outcrossing, making our observation of excess heterozygosity a conservative estimate. We further estimated selfing rates using the robust multilocus estimate of selfing (RMES)^17^, which relies on the distribution of multilocus heterozygosity rather than on fixation indices and is therefore not biased by miscalled homozygous sites in sequence data. Across most populations, RMES confirmed negligible selfing, where the identity disequilibrium (*g*_2_) was not significantly different from zero in Central Europe (Belgium), North America (Massachusetts), South America (Bariloche), and both Australian populations (all *p* > 0.1, 1,000 permutations; Table 2). Population-level statistics could not be calculated for Africa because only one individual was available. Only New Zealand had a statistically significant *g*_2_, corresponding to an estimated selfing rate of 5.60% (*p* = 0.004). Maximum-likelihood estimation gave a consistent value of 6.85% in New Zealand and zero in all other populations. The elevated selfing rate in New Zealand was driven entirely by three individuals of recent selfing origin (NZ105, NZ106, and NZ174; see below) rather than by a population-wide shift. When these three individuals were excluded, *g*_2_ was no longer significant and the maximum-likelihood selfing rate dropped to zero. Outcrossing therefore predominates across native and introduced populations alike.

**Table 2.**
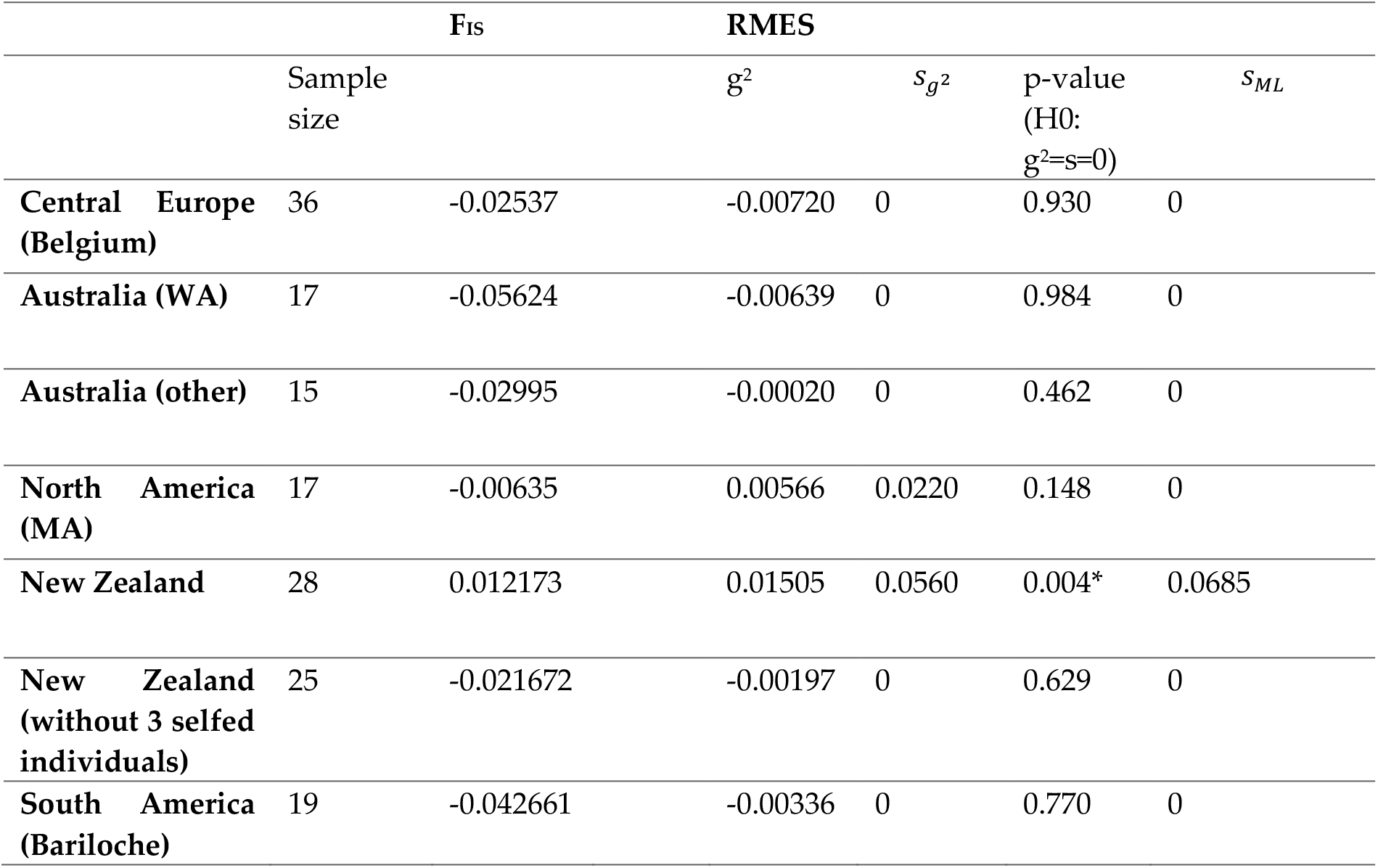
Per-population sample sizes, inbreeding coefficients, and selfing rate estimates. *F*_IS_, Nei’s inbreeding coefficient. *g*_2_, second-order identity disequilibrium. *S*_g2_, selfing rate from *g*_2_. *S*_ML_, maximum-likelihood selfing rate.

### Three New Zealand individuals are products of direct selfing

Individual-level inbreeding analysis traced the three New Zealand outliers as the source of the elevated population-level selfing signal (Figure 2). NZ105 and NZ174 had approximately half the heterozygosity of other New Zealand individuals and inbreeding coefficients (*F*_ROH_, *F*_hom_) near 0.5, consistent with being direct offspring of mating between siblings from the same parent. NZ106 had approximately three-quarters of the average heterozygosity, suggesting it may be the result of a cross between a selfed individual and its half-sibling. All three individuals were collected from isolated pine hosts in urban or semi-urban parks rather than in dedicated pine plantations. Notably, other individuals collected from comparable park or horticultural environments, including those in North America, did not show this signature, indicating that selfing remains rare even in low-density park habitats. The only available sporocarp from Africa also showed inbreeding coefficients near 0.5 and a fraction of heterozygous sites comparable to NZ105 and NZ174, suggesting that it likely also had a selfing origin, although the population-level selfing rate cannot be estimated. In all four individuals, allele frequencies at heterozygous sites were centered at 0.5, confirming single-peak distributions and excluding cross-contamination or polyploidy as artifact sources^12^. Outside these four outliers, the distribution of inbreeding coefficients and heterozygosity were continuous within every population, with no evidence of additional recent selfing events. To our knowledge, NZ105 and NZ174 represent the first cases of individuals arising through selfing detected in any bolete.

**Figure 2.**
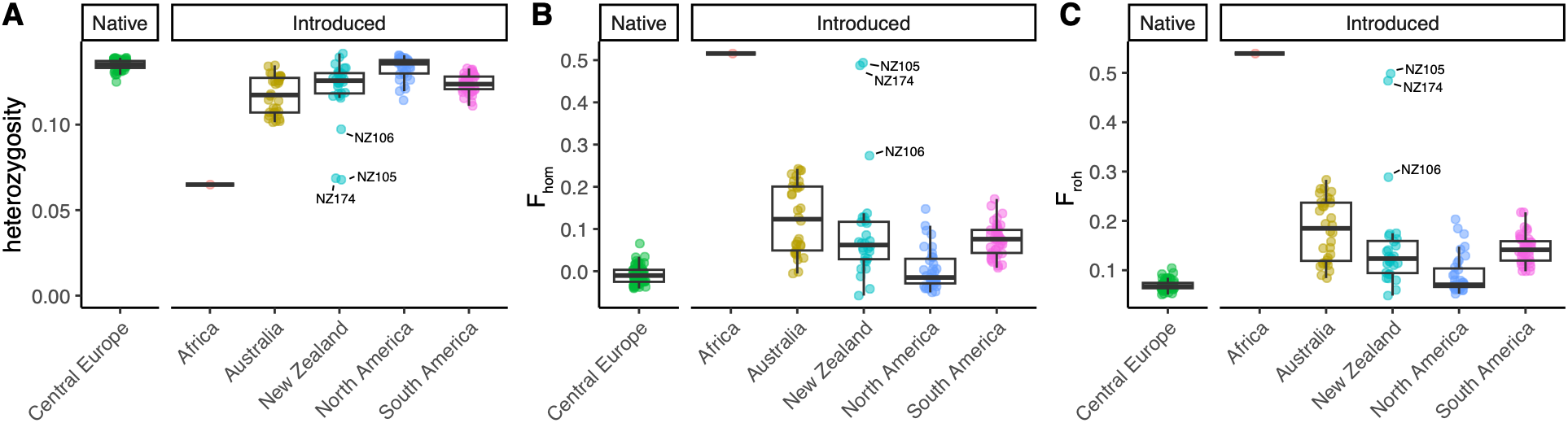
Three New Zealand individuals show inbreeding signatures consistent with recent diploid selfing. Per-individual estimates of inbreeding across native and introduced populations of *S. luteus*. Each point depicts (A) the proportion of heterozygous sites, (B) excess homozygosity (*F*_hom_), and (C) inbreeding coefficient from runs of homozygosity (*F*_ROH_) for an individual. Three New Zealand individuals (NZ105, NZ106, NZ174) deviate sharply from their population background. NZ105 and NZ174 have approximately half the heterozygosity of other New Zealand individuals and *F*_hom_ and *F*_ROH_ near 0.5, consistent with direct sibling-mated offspring of a single parent. The single African sporocarp shows a similar pattern. NZ106 has approximately three-quarters the normal heterozygosity, suggesting a cross between a selfed individual and a half-sibling.

### Inbreeding depression, not mating-system architecture, is the barrier to selfing in S. luteus

Our results suggest that selfing is selectively disfavored across both the native and introduced range of *S. luteus* despite being permitted by the breeding system. If sustained mixed mating were occurring, a continuous distribution of heterozygosity together with a significant population-level selfing rate from RMES would be expected. In a bipolar basidiomycete with a single-locus SI system, such as *S. luteus*, half of the monokaryotic meiotic products from a single dikaryotic parent are expected to be compatible with their siblings at the *MAT* locus^10^. The near absence of selfing must therefore be enforced by other mechanisms, of which we interpret inbreeding depression as the most likely primary barrier. Inbreeding depression has been observed to result in a 15-24% reduction in fitness across both cultivated and wild basidiomycetes^18–20^. Comparable costs in ascomycete fungi^21,22^, suggest that inbreeding depression is a widespread feature across the fungal tree of life. Although a direct test of inbreeding depression is lacking in *Suillus*, we observed that the inbreeding load^23^ (the number of heterozygous loss-of-function variants) in *S. luteus* was 1.3 to 1.8 times higher than that in *A. phalloides* (Fig. 3). Furthermore, relatively lower ratio of expressed load^23^ (the number of homozygotes variants) to inbreeding load of loss-of-function variants were detected in *S. luteus* but not in *A. phalloides* when compared to neutral sites, implying the loss-of-function variants are more strongly selected against in *S. luteus* populations. If the high outcrossing rate in *S. luteus* is maintained by inbreeding depression driven by deleterious recessive alleles, as in most outcrossing systems, the species likely harbors substantial genetic load. Both selfing and unisexual reproduction would expose these deleterious alleles in the dominant dikaryotic stage of the fungus’s life cycle. This cost could explain why neither selfed nor unisexual lineages have established in introduced *S. luteus* populations, despite the advantage predicted by Baker’s rule.

**Figure 3.**
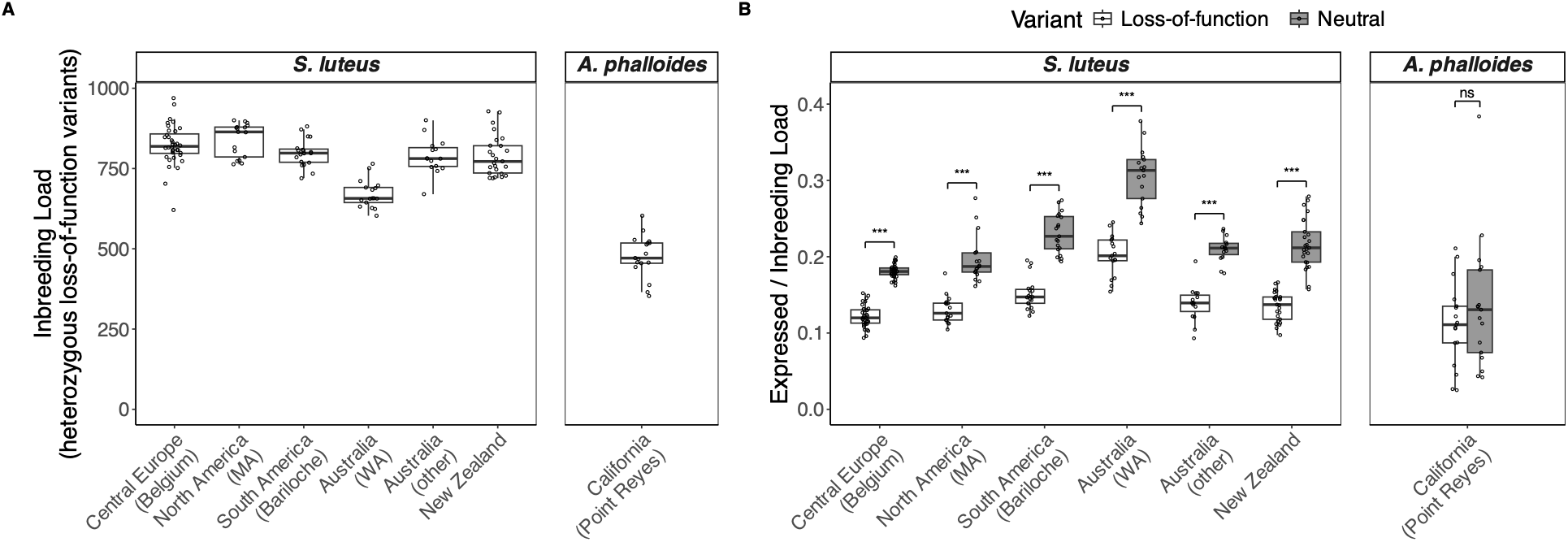
Inbreeding load of loss-of-function variants per individual in populations of *Suillus luteus* and *Amanita phalloides*. (A) The inbreeding load as measured by the number of heterozygous loss-of-function variants (disruption of start and stop codons) per individual (B) Ratio of expressed load (alleles in homozygotes) to inbreeding load for loss-of-function variants versus neutral (synonymous) variants per individual. Variants were polarized with the minor allele as derived. Each point is an individual. Boxes show the median and interquartile range. Brackets show two-sided sign tests comparing loss-of-function and neutral variants within individuals (*** p < 0.001; * p < 0.05; ns, not significant).

### Challenges in studying fungal mating and breeding systems in natural environments

In plants, selfing rates can be measured directly via progeny array analysis, where seeds collected from a maternal plant of a known genotype are individually genotyped to distinguish selfed versus outcrossed offspring^24^. This approach is not feasible in basidiomycetes because the parental dikaryon (the sporocarp) sheds monokaryotic meiotic spores that must encounter a compatible mate before re-establishing as a dikaryotic offspring. Because the parental and offspring dikaryons are never co-located on the same individual, mating outcomes can only be inferred indirectly from the genotypes of independently sampled dikaryons, necessitating reliance on population-genetic estimates of selfing.

Some boletes, including *Suillus* and *Rhizopogon*, are known to have the developmental potential for pseudo-homothallism^25,26^, in which two meiotic nuclei are packaged into a single spore that germinates as a dikaryotic mycelium, analogous to autogamy in plants. From a genetic perspective, pseudo-homothallism cannot be distinguished from orthodox diploid selfing, in which two monokaryotic spores from the same parent germinate independently before fusing to form a dikaryotic mycelium^10^. Direct observation of spore germination and mating is the only definitive way to distinguish between the two, but is not feasible in natural populations given the microscopic scale at which these processes occur. We note, however, that the downstream selective consequences remain the same and that this distinction does not affect our central inference.

### Two basidiomycete colonizers, two reproductive responses on demographic grounds

The contrast between *S. luteus* and *A. phalloides* highlights two distinct reproductive responses among invasive basidiomycetes. In California, *A. phalloides* produces monokaryotic sporocarps from monokaryotic mycelia in the absence of compatible mates, a developmental innovation that bypasses *MAT* control^7^. Monokaryotic genets of *A. phalloides* have been reported to persist for years without being eliminated via competition^8^. In contrast, *S. luteus* shows neither monokaryotic fruiting nor elevated selfing, maintaining predominant outcrossing even in introduced populations. Several ecological and genetic differences between the two systems may explain the contrasting outcomes. First, the population densities after establishment differ. *S. luteus* is abundant in pine plantations where it can dominate the ectomycorrhizal community and colonize the majority of root tips^27^, providing ample mating opportunities but also stronger intra-specific competition. *A. phalloides* in California occurs in scattered urban and peri-urban habitats under oaks and other broadleaf hosts^28^, where its founder population density and connectivity may be lower. These differences in population density and spatial connectivity may influence the availability of compatible mates and the strength of intra-specific competition, and therefore lead to different consequences in the tradeoff between alternative reproductive strategies and inbreeding depression. Second, the two species may differ in phylogenetic constraint. Unisexual reproduction is phylogenetically patchy across basidiomycetes, with most documented cases in *Cryptococcus*^29^ and *Amanita*^7^, and *S. luteus* may simply lack the capacity for unisexual reproduction. Even so, our data show that the selection does not favor selfing in introduced populations under the existing breeding system, and is therefore unlikely to favor a breakdown of SI system that would further encourage selfing. The barrier to selfing in *S. luteus* may therefore be selective, rather than architectural.

### Implications for Baker’s rule and mating-system evolution in basidiomycetes

Our findings refine the scope of Baker’s rule across kingdoms. In plants, the rule has its strongest empirical support^30,31^, although many successful plant invaders remain obligate outcrossers^2,4^. Pannell et al.^4^ reformulated Baker’s law as predicting an enrichment of the capacity for uniparental reproduction in colonizing populations, rather than an increase in the realized selfing rate. This narrower formulation is most applicable to basidiomycetes, given that unisexual reproduction evolved in *A. phalloides* whereas the selfing rate of *S. luteus* remained low across its introduced range. The differing reproductive responses of *S. luteus* and *A. phalloides* to introduction raise broader questions for basidiomycete invasion biology including whether these responses vary predictably with breeding-system flexibility, the strength of inbreeding depression, population density, and competitive context. Resolving these questions will require wider population-level mating system and breeding system data from more introduced basidiomycete species with varied life-history traits, and dedicated measurement of inbreeding depression. The mating system and breeding system are not merely reproductive features, but affect how genetic variation is generated and maintained, which in turn determine how well introduced fungi adapt to novel environments^9,10,30,32–34^. Building a comparative picture of mating system diversity across the basidiomycete tree of life is a necessary step toward understanding how reproductive strategy shapes the evolutionary trajectories of fungi in the Anthropocene.

## Materials and Methods

### SNP discovery

All samples are genomes from previously published collections^12^ and the same set of SNPs was used. Briefly, reads were trimmed using Cutadapt v2.1^35^ (minimum quality 25; minimum length 80 bp; Illumina TruSeq adaptors) and mapped to the *S. luteus* reference genome UH-Slu-Lm8-n1 v3.0^36^ using BWA v0.7.17^37^. Variant calling was performed using GATK HaplotypeCaller and GenotypeGVCFs^38^. Hard filtering following GATK best practices (QD < 2.0 || FS > 60.0 || MQ < 40.0 || MQRankSum < -12.5 || ReadPosRankSum < -8.0) and DP > 1.5× genome median retained 3,381,371 SNPs from 189 dikaryotic genomes.

### Selfing rate estimation

Population-level selfing rates were estimated by Nei’s F_IS_^39^ using the *hierfstat* package (https://doi.org/10.32614/CRAN.package.hierfstat), and by the robust multilocus estimate of selfing (RMES) program^17^. RMES uses the distribution of multilocus heterozygosity to estimate selfing through identity disequilibrium (g_2_) and maximum-likelihood estimation (ML). Both were computed with 200 unlinked random SNPs after LD-pruning using PLINK v1.9^40^ (−-indep-pairwise 50 10 0.1). Significance was assessed by 1,000 permutations under H0: g2 = s = 0, as implemented in the RMES program. Individual inbreeding was quantified by excess homozygosity (F_hom_) and proportion of heterozygous sites across sites in a single genome using the --het option in PLINK v1.9, and by runs of homozygosity (ROH) using the roh option in bcftools v1.23^41^ with an assumed recombination rate 30 cM/Mb as in model basidiomycete organisms *Coprinopsis cinerea*^42^ and *Pleurotus ostreatus*^43^. ROH regions > 10,000 bp were retained to calculate F_ROH_.

### Mating type determination

*HD MAT* alleles were locally assembled using the method from Ke et al^11^. WGS reads with at least one paired end mapping to the *HD* locus in the reference were collected and *de novo* assembled with MIRA^44^. Multiple sequence alignment was performed with *MAFFT*^45^ for all alleles together with the annotated *HD MAT* locus from the reference genome to determine the boundaries of genes and CDS (Files S1-S3). Pairwise allele identity was calculated from *MAFFT* alignments of the nucleotide sequences of *HD MAT* and the translated amino acid sequence of the *HD1* and *HD2* protein. Both alignments are available in the Supplementary Files. We grouped alleles across genomes by pairwise sequence identity of the 200-aa N-terminal region of the two *HD* proteins within the locus, the region directly involved in mating specificity^46^. The total number of distinct mating types per population was estimated by the incompatible-pairs method and by maximum-likelihood estimation under multinomial distribution assuming equal mating-type frequencies in the population (Methods S1; Figure S3).

### Loss of function alleles annotation

SNPs of all 17 dikaryotic *A. phalloides* genomes from fruiting bodies collected at Point Reyes National Seashore (PRNS), California, USA, were called using the same methods as *S. luteus* with the reference genome and SRA from Wang et al.^7^ The SNPs of both species were categorized by their impact using *SnpEff*^47^ according to genome annotation by funannotate (https://github.com/nextgenusfs/funannotate). The “HIGH” impact SNPs, including disruption of start codon and stop codons gains, were consider loss-of-function variants. The “LOW” impact SNPs, including synonymous variants, were consider neutral variants.

## Acknowledgments

We thank the Office of Information Technology and Research Computing team at Duke University for maintaining and providing computational resources. Financial support for this work was provided through National Science Foundation Grants DEB 1554181 to R.V., IOS-PBI 2029168 to S.B., DEB 2332726 to L.L. This material is based upon work while S.B. was serving at the National Science Foundation. We thank Timothy Y. James and Nhu H. Nguyen for providing valuable feedback on the manuscript.

## Data and code availability

Analysis scripts are available at https://github.com/KeFungi/Suilu_Self.

## Author Contributions

Y.-H.K. and R.V. conceived the study. Y.-H.K. performed analyses and wrote the manuscript with input from all authors. Y.-W.W., A.L.B., L.L., S.B., and R.V. reviewed the analyses and improved the manuscript.

## Supplementary Information

**Figure S1.**
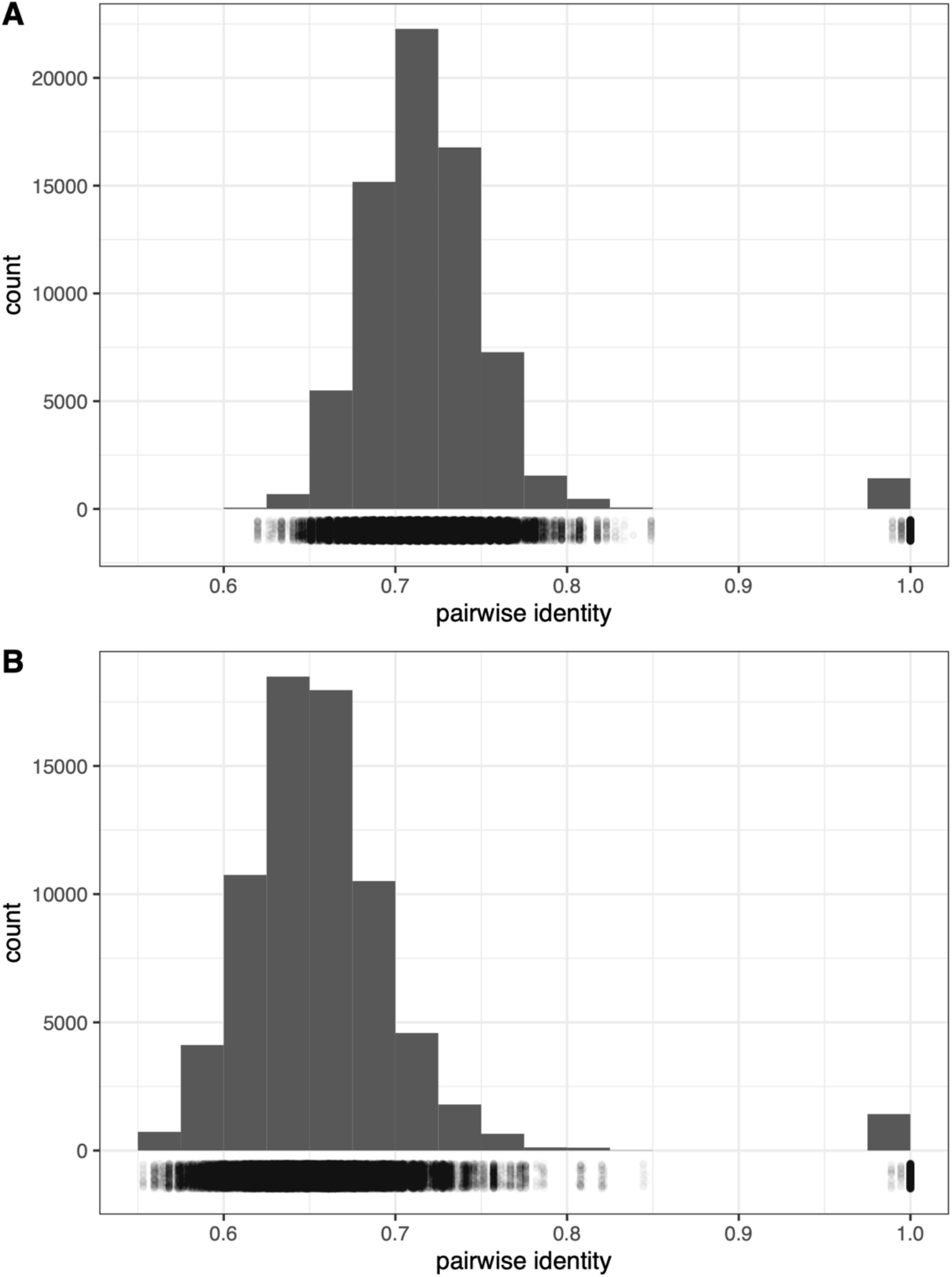
Pairwise sequence identity of *HD MAT* alleles. Distribution of pairwise sequence identities for the 200-aa N-terminal region of assembled (A) *HD1* alleles and (B) *HD2* alleles, across the 189 *S. luteus* individuals. Each dot below the histogram indicates the exact identity value for each pair.

**Figure S2.**
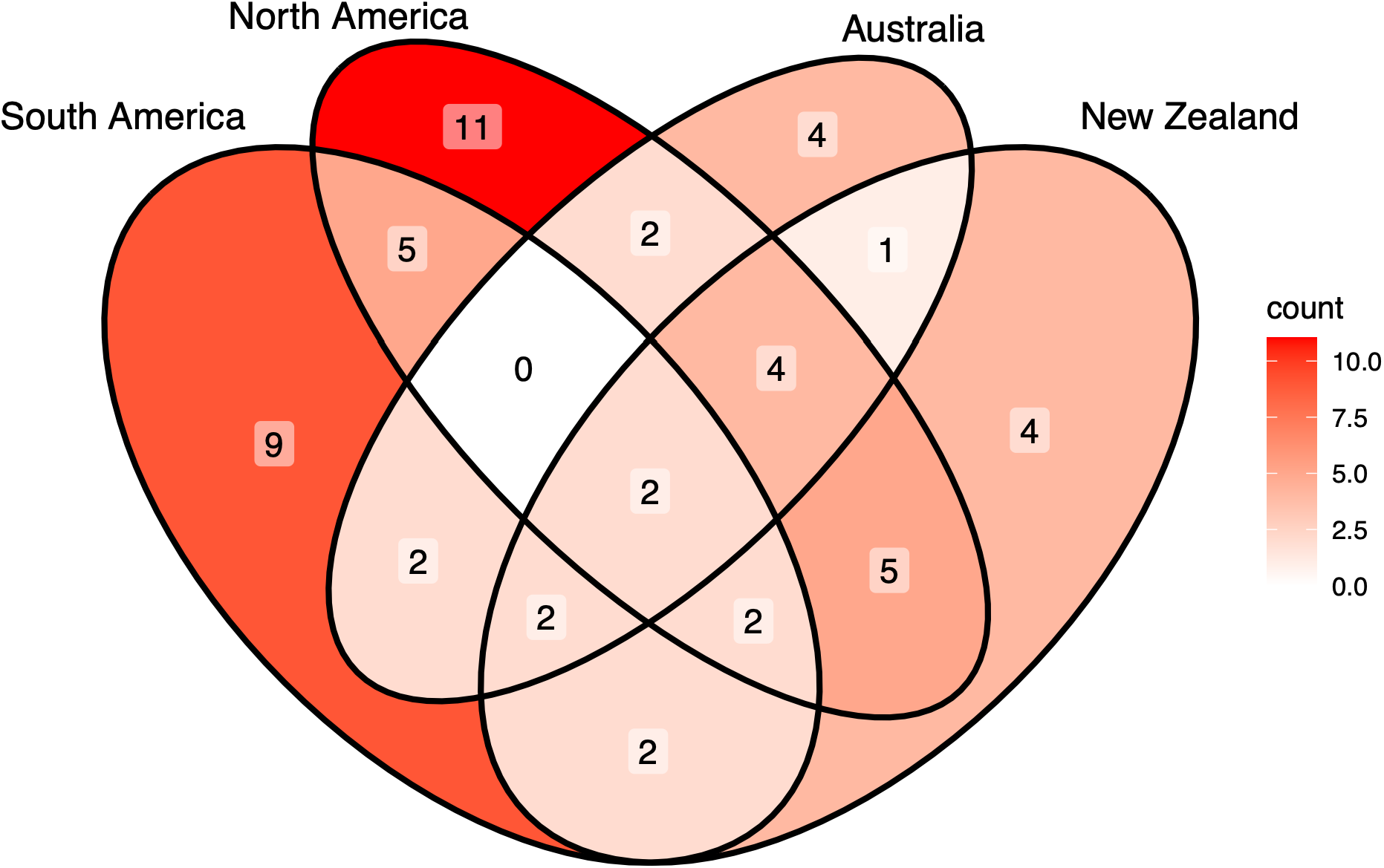
*HD MAT* alleles shared among the four introduced populations. Venn diagram of *HD MAT* alleles shared among introduced populations in North America, South America, Australia, and New Zealand.

**Figure S3.**
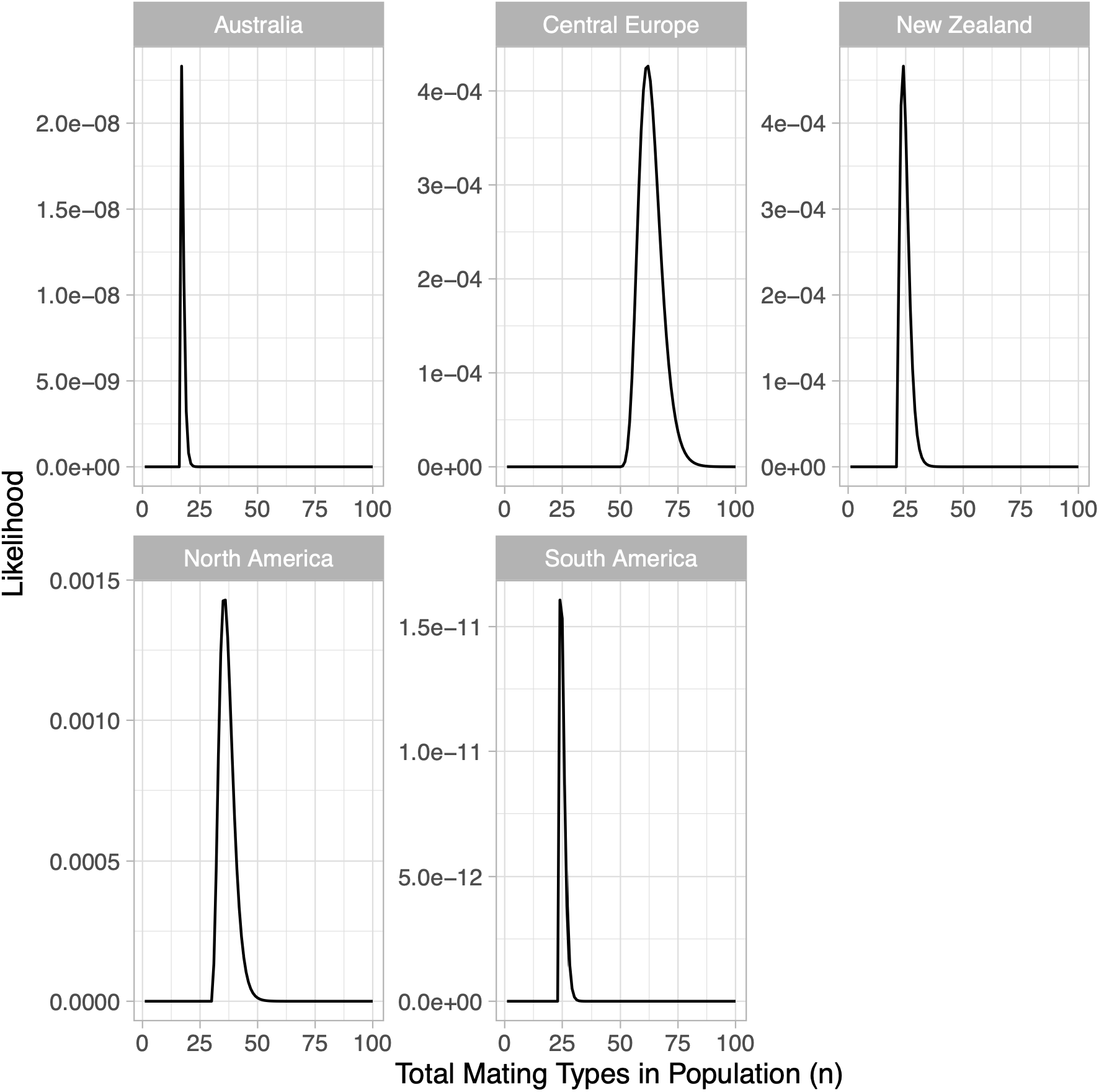
Likelihood surface of total mating types in each population.

**File S1 Alignment of HD MAT locus of all assembled alleles**

**File S2 Alignment of HD1 protein of all assembled alleles**

**File S3 Alignment of HD2 protein of all assembled alleles**

**Table S1. Pairwise sequence identity matrix of N-terminal 200-amino-acid sequences of *HD1* alleles.**

**Table S2. Pairwise sequence identity matrix of N-terminal 200-amino-acid sequences of *HD2* alleles.**

**Table S3. Per-individual *HD MAT* mating type assignments.**

**Table S4. *HD MAT* allele counts and per-population estimates.**

## Supplementary Methods

### Estimate number of mating types in populations

Raper, Krongelb, and Baxter (1958) estimated the total number of mating types in a fungal population by the number of incompatible pairs, a concept originated from the study of the frequency of lethal recessive alleles in *Drosophila* (Dobzhansky and Wright 1941). It assumes a population has a total of *n* mating types of equal frequencies 1/*n*. Therefore, the probability to sample two alleles of the same type is 1/*n*. In the context of crossing, let the number of incompatible pairs of gametes *d*. The expected ratio of incompatible pairs of gametes in all combinations of *k* tested gamete equals 1/*n*

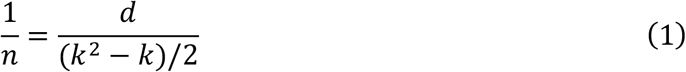

In this case, since the mating types have been assigned from sequences, the results of crossing can be predicted. Assuming allele *A*_*i*_ is sampled *k*_*i*_ times, we have the number of incompatible pairs from *A*_*i*_

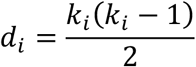

The *n* can be obtained from (1) according to

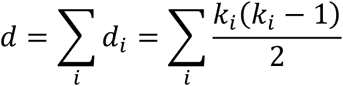

and *k* from

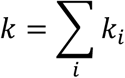

in this putative crossing.

The number of mating types in a population can also be estimated from the exact distribution of *k*_*i*_ based on the multinomial distribution. Using the same assumption that all *n* types of alleles have the same frequency, *p*_1_ = *p*_2_ = ⋯ = *p*_n_ = 1/*n*, the probability to sample *A*_1_ *k*_1_ times, *A*_2_ *k*_2_ times, …, *A*_n_*k*_n_ times following multinomial distribution is

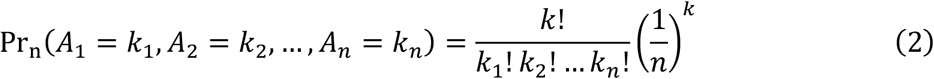

Since the numbering of allele types is arbitrary, the order of observed allele counts is exchangeable. For example, the probability of three allele types sampled *k*_1_, *k*_2_, *k*_3_ times, with k_1_, k_2_, k_3_ pairwise distinct, in a population of 3 allele types is

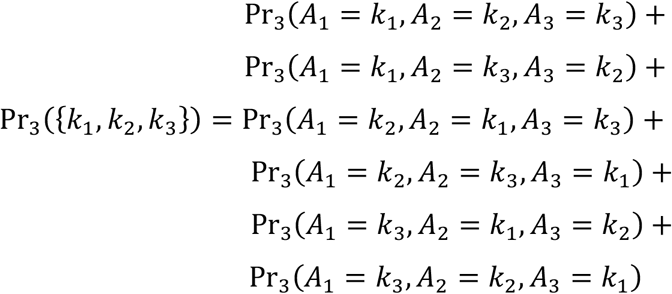

Because the probability for each case is equal, it is equivalent to the counts for each allele type in any order multiplying the number of permutations of {*k*_1_, *k*_2_, *k*_3_}

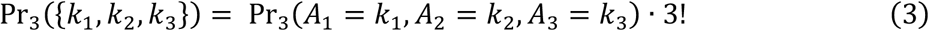

However, the counts of different allele types are not always pairwise distinct. When some k_i_ values are equal, the factor 3! in (3) generalizes to P, the number of distinct orderings of the multiset {k_1_, k_2_, …, k_n_}

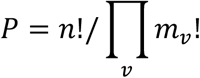

where *m*_*v*_ = #{*i*: *k*_*i*_ = *v*} is the multiplicity of count value v among k_1_, …, k_n_ (including v = 0 when fewer than n allele types are observed). Then,

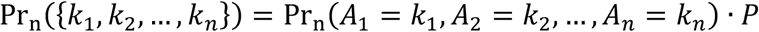

The likelihood function for the number of allele types n given observed counts {k_1_, …, k_n_}, padded with zeros if fewer than n types are sampled, is

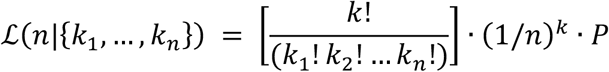

where P is defined as above. The likelihood is zero when the number of observed allele types exceeds n. For example, when n = 5 and k_1_, k_2_, k_3_ are pairwise distinct, the observed sample in (3) becomes

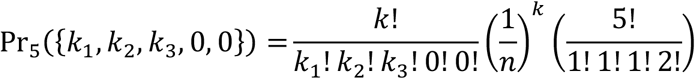

Using this likelihood function, maximum-likelihood estimation for the number of allele types in population (*n*) was performed for *n* from 1 to 100.

